# Tissue-bound hyaluronan molecular weight as a regulator of dendritic cell immune potency

**DOI:** 10.1101/2023.06.06.543835

**Authors:** Brian Chesney Quartey, Jiranuwat Sapudom, Mei ElGindi, Aseel Alatoom, Jeremy Teo

**Affiliations:** Laboratory for Immuno Bioengineering Research and Applications, Division of Engineering, New York University Abu Dhabi, Abu Dhabi, UAE; Department of Mechanical Engineering, Tandon School of Engineering, New York University, USA; Department of Biomedical Engineering, Tandon School of Engineering, New York University, USA

**Keywords:** Dendritic cells, innate immune response, hyaluronan, molecular weight, CD44, extracellular matrix

## Abstract

Hyaluronic acid (HA) is a major glycosaminoglycan found in the extracellular matrix (ECM) and exhibits immunoregulatory properties depending on its molecular weight (MW). However, the impact of tissue bound HA on dendritic cell (DC) functions is not well understood due to the varying distribution of HA MW under different physiological and pathological conditions. To investigate DCs in defined biosystems, we used three-dimensional (3D) collagen matrices modified with HA of specific MW, while maintaining similar microstructure and HA levels. Using these matrices, we examined the influence of HA on cytokine binding and observed distinct properties depending on the presence and MW of HA, suggesting modulation of cytokine availability by the different MW of HA. Our studies on DC immune potency revealed that low molecular weight HA (LMW-HA; 8-15 kDa) enhances immature DC (iDC) differentiation and antigen uptake, while medium (MMW-HA; 500-750 kDa) and high molecular weight HA (HMW-HA; 1250-1500 kDa) increase cytokine secretion in matured DCs (mDCs). Interestingly, the modulation of DCs surface marker expression and cytokine secretion by different MW of HA appeared to be independent of CD44. However, we found that cytokine secretion of DCs was dependent on the CD44 receptor regardless of the presence or absence of HA in the matrix. Additionally, we observed reduced migratory capacity of iDCs and mDCs when cultured on MMW- and HMW-HA matrices, and this effect was dependent on CD44. In summary, our findings provide new insights into the MW-dependent effects of tissue-bound HA on DCs, opening avenues for the design of DC-modulating materials to enhance DC-based therapy.

## 1. Introduction

Dendritic cells (DCs) are the most potent antigen-presenting cells of the innate immune system and play a crucial role in educating the adaptive immune response [1]. They exist in two distinct functional and phenotypic states, namely immature and mature[2]. In the immature state, dendritic cells (iDCs) reside in non-lymphoid tissues where they scavenge for foreign antigens. iDCs recognize foreign antigens through specialized pattern recognition receptors and phagocytose them[1]. Following this, iDCs undergo maturation, transforming into mature DCs (mDCs). mDCs then migrate from peripheral tissues to lymphoid organs and present antigens on major histocompatibility complex II (MHC II) proteins to T cells, thereby activating them and initiating the adaptive immune response[3,4]. Biomechanical and biochemical cues inherent in tissues can modulate the immunological functions of DCs as they scavenge peripheral tissues and migrate through different tissues[5]. Previous studies have reported that matrix stiffness[6,7] and tissue density[8] can influence DC functions and the maturation process. However, the impact of various extracellular matrix (ECM) components on DC immune potency remains poorly understood.

One crucial ECM component, hyaluronan (HA), is a glycosaminoglycan that exhibits a high turnover rate and possesses immunomodulatory properties[9–11]. HA is known to impact hydration, lubrication, and physical properties of tissues[12,13]. It consists of a repeating disaccharide chain composed of [3)-β-d-N-acetylglucosamine (GlcNAc)-β(1,4)-d-glucuronic acid (GlcA)-β(1)][14]. HA contributes to the structural integrity of the ECM and triggers various signaling pathways, regulating cellular processes through its interaction with principal receptors, CD44 and RHAMM[14]. Notably, HA exhibits contradicting roles in different biological processes depending on its physical properties, such as molecular weight (MW) distribution, chemical modifications, metabolism, and interactions with receptors, particularly CD44[15]. In vivo, HA in the ECM exists in different molecular weight ranges depending on various physiological contexts, ranging from high MW (HMW-HA) of 1000–8000 kDa to low MW (LMW-HA) of 50 kDa or less[12,16]. Three main integral proteins called HA synthases (HAS1, HAS2, HAS3) are involved in synthesizing different sizes of HA[17,18]. HA is synthesized inside the cell membrane of fibroblasts and extruded into the extracellular space[19]. Under normal physiological conditions, HA is synthesized in its HMW form. However, during certain physiological and pathological conditions, such as tissue injury and cancer microenvironments, HMW-HA can be broken down into smaller fragments of different sizes through depolymerization by reactive oxygen species or enzymatic degradation by hyaluronidases (HYAL)[20]. Several studies have demonstrated that HAs of different MW distinctly regulate immune responses. Recent research has shown that LMW-HA upregulates pro-inflammatory genes, including nos2, TNFα, IL-12β, and CD80, and enhances secretion of nitric oxide and TNFα in macrophages[21]. On the other hand, HMW-HA upregulates anti-inflammatory genes, including arg1, IL-10, and MRC1[21]. In another study, different MW HA has been shown to modulate macrophage polarization, with HMW-HA inducing polarization of unstimulated macrophages toward a more M2, anti-inflammatory, phenotype[22]. These findings suggest that HMW-HA suppresses the pro-inflammatory response. While these studies shed light on the effects of varying MW HA on macrophage activation and response, less is known about the impact of different MW HA on DCs within physiologically relevant tissue microenvironments. Understanding how HA instructs DC phenotypes and functions will enhance current approaches for immunoengineering DCs and pave the way for novel therapies.

Given the challenges in deciphering the impact of HA MW in regulating dendritic cell immune potency in native tissues, this study aims to utilize biomimetic three-dimensional (3D) collagen matrices with immobilized HA of well-defined MW. The differentiation and maturation of dendritic cells cultured within these collagen matrices were investigated through cell surface marker expression, cytokine secretion profiles, antigen uptake capabilities, and migration assays, with a focus on their dependency on the CD44 receptor.

## 2. Materials and Methods

### 2.1. Reconstruction of 3D collagen matrices

Collagen matrices were prepared using a well-established protocol, as previously published[23]. In brief, rat tail type I collagen (Advanced BioMatrix, Inc., CA, USA) was mixed with 0.1% acetic acid (Merck KGaA, Germany) and 0.5 M phosphate buffer (Merck KGaA, Munich, Germany) on ice to achieve a collagen concentration of 2 mg/ml. The prepared collagen solution was then transferred onto glutaraldehyde-coated coverslips (13 mm in diameter; VWR, Leuven, Germany) and placed under standard cell culture conditions of 37°C, 5% CO_2_, and 95% humidity to allow collagen fibrillation. Following fibrillation, the collagen matrices were washed and stored in phosphate-buffered saline (PBS; Thermo Fisher Scientific Inc, Leicestershire, UK) until further use.

### 2.2. Chemical crosslinking of 3D collagen matrices

To crosslink the reconstituted collagen matrices, they were incubated with 4 mg/mL of N-(3-Dimethylaminopropyl)-N′-ethylcarbodiimide hydrochloride (EDC; Sigma-Aldrich, Schnelldorf, Germany) dissolved in 100 mM 2-(N-morpholino) ethanesulfonic acid (MES; Sigma-Aldrich, Schnelldorf, Germany) at pH 5 for 2 hours at room temperature, following a previously published method[24]. After crosslinking, the matrices were washed three times with PBS (Thermo Fisher Scientific Inc, Leicestershire, UK) and then incubated with cell culture medium for 24 hours before use.

### 2.3. HA immobilization into 3D collagen matrices

The immobilization of HA onto reconstituted 3D collagen matrices was carried out following a previously established protocol[18,20]. To immobilize a defined amount of 1 μg of HA per matrix, the following concentrations of HA were dissolved in 100 mM MES buffer (Sigma-Aldrich, Schnelldorf, Germany) at pH 5: 9.1 µg/ml of LMW-HA (8-15 kDa), 13.5 µg/ml of MMW-HA, (500-750 kDa), and 0.02 µg/ml of HMW-HA (1250-1500 kDa) (Contipro Biotech. S.r.o., Dolni Dobrouc, Czech Republic). The matrices were then incubated with the prepared HA solutions for 2 hours at room temperature. Subsequently, the HA solution was removed without a washing step, and a prepared crosslinking solution of 4 mg/mL EDC (Sigma-Aldrich, Schnelldorf, Germany) in MES buffer at pH 5 (Sigma-Aldrich, Schnelldorf, Germany) was directly added to the matrices and incubated for 2 hours at room temperature. The EDC crosslinker enables the covalent immobilization of HA to the 3D collagen matrices[25]. Following crosslinking, the matrices were washed three times with PBS (Thermo Fisher Scientific Inc, Leicestershire, UK) and incubated with cell culture medium for 24 hours before use.

### 2.4. Quantification of immobilized HA using Alcian blue assay

The amount of HA immobilized on 3D collagen matrices was quantified following a previously published protocol[18]. In brief, the 3D collagen matrices were digested using a 6 mg/ml solution of type IV collagenase (Worthington Biochemical Corporation, USA). The digested collagen matrices were then incubated with Alcian blue 8GX (Sigma-Aldrich, Schnelldorf, Germany) for 30 minutes at room temperature while being placed on a shaker. Alcian blue 8GX reacts with HA, forming an insoluble HA-Alcian blue complex, which was isolated by centrifugation at 15,000g for 15 minutes at 4°C. The sedimented complex was subsequently washed twice with deionized water. Next, the formation of the HA-Alcian blue complex was reversed by adding 8M guanidine hydrochloride (dissolved in deionized water) (Sigma-Aldrich, Schnelldorf, Germany). The dissolved Alcian blue, dependent on HA, was quantified by measuring the absorption at 620 nm using a UV/Vis-spectrophotometer (Cytation 5 imaging reader: BioTek, Winooski, VT, USA). A standard curve was used to correlate the absorption value with the amount of HA. The experiments were performed in six replicates.

### 2.5. Characterization of topological and mechanical properties of reconstituted 3D matrices

For topological analysis, 3D collagen matrix systems were stained with 50 µM of 5-(and-6)-carboxytetramethylrhodamine succinimidyl ester (TAMRA-SE, Sigma-Aldrich) at room temperature for 60 minutes and visualized using a confocal laser scanning microscope (cLSM) (SP8; Leica, Germany) equipped with a 40× water immersion objective (Leica, Germany). The acquired cLSM stacked images were subsequently analyzed using a custom-built MATLAB script (MATLAB 2022b; MathWorks, USA), as described in a previous publication[26]. To quantify the pore size of the collagen matrices, stacked images obtained from four different random positions per matrix were analyzed for each of the four reconstituted matrices. The mechanical properties of the 3D matrix systems were analyzed using a non-destructive rheological measurement method employing ElastoSensTM Bio (Rheolution, Quebec, Canada), as outlined in a previous publication[8]. All experiments were performed in four replicates.

### 2.6 Quantification of cytokine binding properties of reconstituted 3D matrices

To investigate the cytokine binding properties of the 3D matrices, we incubated 10 µg/ml of each cytokine (IL-6, IFNγ, CCL2, IL-1β, IL-12p70, IL-2, CXCL8, IL-17A, IL-10, TGF-β1, IL-4, and CXCL10) (BioLegend, San Diego, CA, USA) with the respective matrix conditions for 24 hours under standard cell culture conditions. After incubation, we collected the supernatants and quantified the concentrations of each cytokine using the LEGENDplex™ Human Essential Immune Response Panel (13-plex) (BioLegend, San Diego, CA, USA), following the manufacturer’s instructions. The samples were analyzed using a Cytek Aurora flow cytometer (4L 16V-14B-10YG-8R). Subsequently, quantitative analysis was performed using LEGENDplex™ Data Analysis Software version 8.0 (VigeneTech, USA). To determine the amount of each cytokine bound to the matrix conditions in picograms per microgram of matrix, we subtracted the concentrations of cytokines initially incubated from the concentrations present in the supernatant. All experiments were conducted in six replicates.

### 2.7 Cell culture and differentiation of dendritic cells

The human monocytic THP-1 cell line (AddexBio, San Diego, CA, USA) was maintained in RPMI-1640 cell culture media supplemented with 10% fetal bovine serum (FBS), 1% penicillin/streptomycin, 1% 4-(2-hydroxyethyl)-1-piperazineethanesulfonic acid (HEPES), 1% sodium pyruvate, and 0.01% beta-mercaptoethanol under standard cell culture conditions of 37°C, 5% CO2, and 95% humidity. All cell culture reagents were purchased from Invitrogen, CA, USA.

For cell studies, 5 × 10^4^ THP-1 cells were seeded onto prepared collagen matrices. Differentiation of THP-1 into immature (iDC) and matured (mDC) dendritic cells was performed according to a previously established protocol [8]. Briefly, to differentiate THP-1 cells into iDCs, cells were cultured for 3 days in FBS-free RPMI-1640 cell culture media supplemented with 200 ng/ml IL-4 and 100 ng/ml GM-CSF. For differentiation into mDCs, THP-1 cells were cultured for 3 days in FBS-free RPMI-1640 cell culture media supplemented with 200 ng/ml IL-4, 100 ng/ml GM-CSF, 20 ng/ml TNFα, and 200 ng/ml ionomycin (Sigma-Aldrich, Germany). All cytokines for DC differentiation were purchased from Biolegend, San Diego, CA, USA.

### 2.8. Blocking of CD44 receptor

THP-1 cells were incubated with 2 µg/ml of anti-CD44 antibodies (clone: C44Mab-5; Biolegend, San Diego, CA, USA) for 20 minutes under standard cell culture conditions (95% humidity and 5% CO2 at 37°C) prior to iDC and mDC differentiation. Additionally, 1 µg/ml of anti-CD44 antibodies were added to both activation media. Successful blocking was confirmed by staining with anti-human CD44 antibody conjugated with Alexa Fluor 594 (clone: C44 Mab-5; Biolegend, USA) and quantified using the Cytek Aurora flow cytometer (4L 16V-14B-10YG-8R) (Supplementary Figure S1).

### 2.9. Quantitative analysis of cell surface markers and cell viability using flow cytometry

To evaluate the expressions of cell surface markers, matrices were digested using a 6 mg/ml solution of type IV collagenase (Worthington Biochemical Corporation, USA) for 20 minutes under standard cell culture conditions. The prepared collagenase solution was supplemented with Human TruStain FcX™-Fc receptor blocking solution at a dilution of 1:250. Cells were then stained with mouse anti-human antibodies against CCR7, CD11c, CD44, CD86, CD206 (MMR1), CD209 (DC-SIGN), HLA-DR, and DRAQ5 (viability dye) on ice for 30 minutes. All antibodies, as well as the DRAQ5 dye, were diluted in FBS-free RPMI-1640 cell culture media at a ratio of 1:250. All staining antibodies and the viability dye were purchased from Biolegend, San Diego, CA, USA. Details of all antibodies are provided in Supplementary Table S1. The stained cells were analyzed using the Cytek Aurora flow cytometer (4L 16V-14B-10YG-8R), which utilizes an unmixing (compensation) program from the SpectroFlo software. Data analysis was performed using FlowJo software (Becton, Dickinson and Company, NJ, USA). The experiments were performed in at least 5 replicates.

### 2.10. Quantitative analysis of cytokine secretion profile

Cell culture supernatants were collected after 3 days of dendritic cell (DC) differentiation and maturation. The LEGENDplex™ Human Essential Immune Response Panel (13-plex) (BioLegend, San Diego, CA, USA) was employed to quantify the secreted cytokines in the cell culture supernatant following the manufacturer’s instructions. The samples were analyzed using the Cytek Aurora flow cytometer (4L 16V-14B-10YG-8R). Quantitative analysis was subsequently performed using LEGENDplex™ Data Analysis Software version 8.0 (VigeneTech, USA). At least 5 replicates per condition were analyzed.

### 2.11. Analysis and imaging of ovalbumin uptake by iDC

To analyze the uptake of ovalbumin, immature dendritic cells (iDC) were incubated with FBS-free RPMI-1640 medium supplemented with 10 μg/mL of fluorescein isothiocyanate-conjugated ovalbumin (FITC-ovalbumin; Invitrogen, Germany) for 1 hour under standard cell culture conditions. The 3D matrices were then digested using a solution of type IV collagenase (6 mg/mL; Worthington Biochemical Corporation, USA). Subsequently, the fluorescent signal of the samples was measured using the Cytek Aurora flow cytometer (4L 16V-14B-10YG-8R), which employs an unmixing (compensation) program from the SpectroFlo software. The geometric mean fluorescence intensity was quantified using FlowJo Software version 10.5.3 (BD, USA). The experiments were conducted in at least 5 replicates.

### 2.12. Cell staining, imaging and quantification of cell infiltration into 3D collagen matrices

Both migration of iDc and mDc into 3D matrices was quantified by analyzing the Hoechst-33342 fluorescence signal from individual cell nuclei, as previously described[23,27]. Briefly, cells were fixed with 4% paraformaldehyde (Biolegend, San Diego, CA, USA) for 10 minutes and subsequently permeabilized with 0.1% Triton X100 (Merck KGaA, Darmstadt, Germany) for 10 minutes. Cells were washed with PBS three times after each step. Subsequently, cells were stained with Hoechst-33342 (diluted 1:10,000 in PBS; Invitrogen, Carlsbad, CA, USA). Z-stack images with an interval of 5 µm and an overall z-layer of 500 µm were acquired using a Lionheart FX automated microscope with a 10× objective (BioTek, Winooski, VT, USA). The z-stacked images were used to quantify the percentage of migrating cells and the migration depth using a custom-built MATLAB script (MATLAB R2022b; MathWorks Inc., USA). The percentage of migrating cells was defined as the percentage of cells located more than 10 μm beneath the collagen matrix surface. Maximum migration depth was defined as the distance crossed by 10% of all migrating cells. Four random positions were analyzed per matrix condition in three independent experiments.

### 2.13. Statistical analysis

All experiments were conducted in a minimum of three independent replicates, unless otherwise specified. Error bars indicate the standard deviations (SD). Statistical significance was determined using a Mann-Whitney test, performed using GraphPad Prism 9 (GraphPad Software, USA). The significance level was set at p ≤ 0.05.

## 3. Results and Discussion

While several studies have demonstrated the immunoregulatory properties of various MWs of HA on immune cells such as macrophages and T cells[21,22,28,29], limited information is available regarding the impact of HA and its different MWs on the function of DCs. Furthermore, the existing studies that have partially examined the effect of different MW HA on DCs were conducted using 2D culture systems by supplementing cell culture media with HA[30–32], which does not accurately account for the presentation of tissue-bound HA within the *in vivo* microenvironment to DCs. In our study, we employed a biomimetic tissue niche where HA with varying MWs is immobilized onto 3D collagen matrices, thereby accounting for the dimensionality as well as the presence of tissue bound HA in native tissues. Using these matrices, we investigated the influence of different MW HA on DC differentiation and maturation, antigen uptake, and migratory capacity. **Figure 1A** illustrates the experimental design and setup employed in this study.

**Figure 1:**
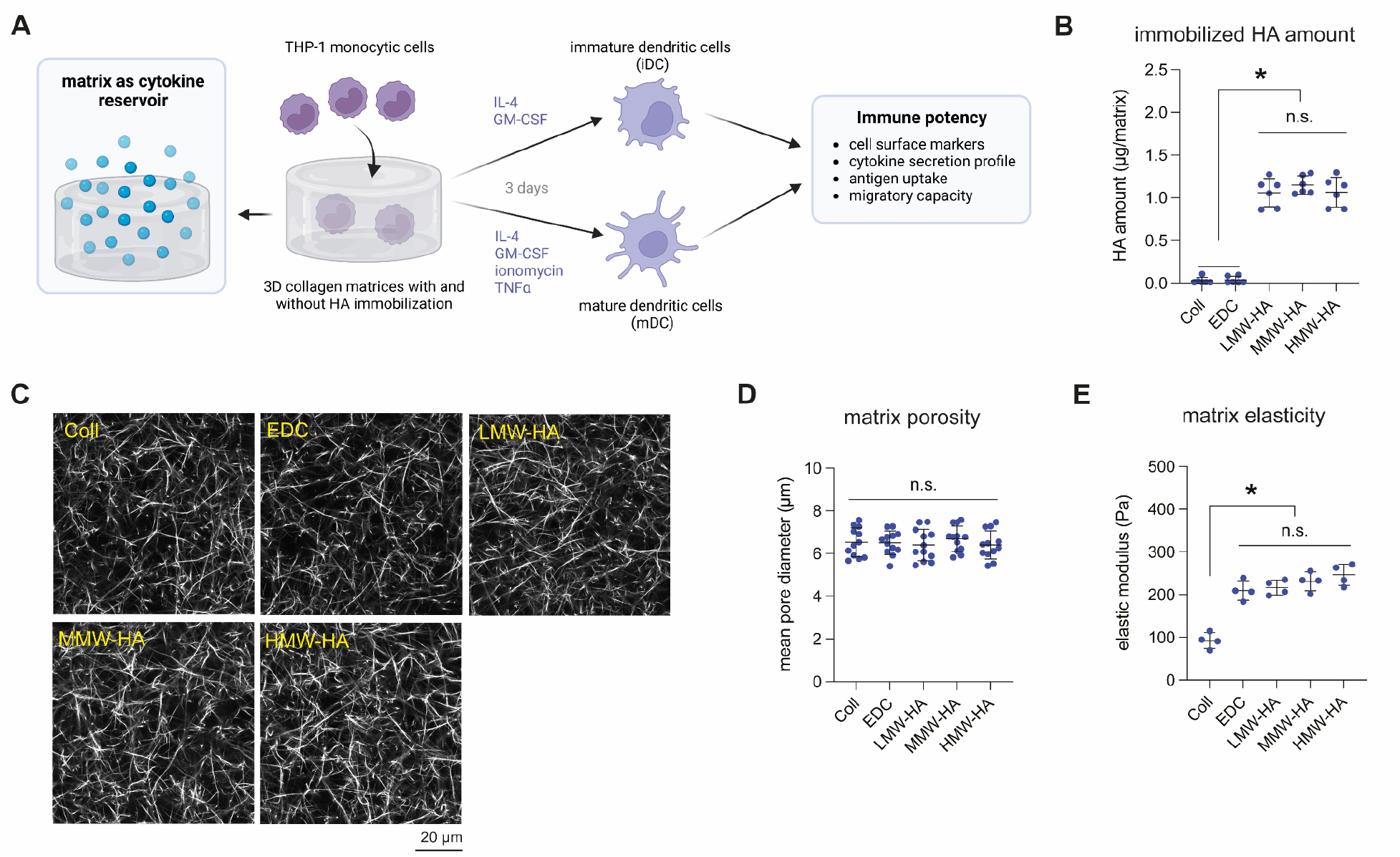
Experimental setup and characterization of reconstituted 3D collagen matrices with and without HA immobilization. (A) Schematic illustration of the experimental setup. HA of varying molecular weights was immobilized onto reconstituted collagen matrices using EDC crosslinking. DCs were differentiated and maturated onto those reconstituted matrices and analyzed with respect to their immune potency. **(B)** The amount of immobilized HA was quantified using the Alcian blue assay (n=6). **(C)** Representative images of collagen matrices with immobilized HA of different molecular weights visualized using confocal laser scanning microscopy (scale bar = 20 µm). **(D)** Matrix porosity was quantified using a custom-made image analysis toolbox. Three different random positions per matrix were analyzed (n=4). **(E)** Matrix elasticity was measured using a non-destructive rheometer (n=4). * and n.s. indicate a significance level of p ≤ 0.05 and non-significant, respectively, as determined by the Mann– Whitney test.

### 3.1. Reconstitution and characterization of 3D collagen matrices with immobilized HA of different MW

Collagen matrices were reconstituted and post-functionalized with hyaluronic acid (HA) of different molecular weights, namely low molecular weight HA (LMW-HA, 8-15 kDa), medium molecular weight HA (MMW-HA, 500-750 kDa), and high molecular weight HA (HMW-HA, 1250-1500 kDa), using EDC chemistry, which enabled covalent binding of adjustable amounts of HA within 3D collagen matrices, as previously described[18,20,25]. Before utilizing the reconstituted matrices for cell studies, we quantified the amount of immobilized HA through an established Alcian blue assay (**Figure 1B**). As expected, only matrices with immobilized HA contained HA, present in comparable amounts of approximately 1 µg per matrix (0.01 µg HA per µg collagen).

Since we have previously demonstrated that the biophysical properties of collagen matrices can regulate the immune potency in DCs[8], we quantified the porosity and elastic modulus of the reconstituted collagen matrices. To visualize the microstructure of the matrices, they were fluorescently labeled and imaged using confocal laser scanning microscopy. **Figure 1C** presents representative images of collagen matrices with and without HA post-modification. It can be visually observed that all matrices exhibited similar microstructure. To confirm this visual observation, we quantified the matrix porosity using a custom-built image analysis toolbox. As depicted in **Figure 1D**, the mean pore diameter remained constant across all matrix conditions, with an average pore size of approximately 6.5 µm. In addition to matrix porosity, the mechanical properties of the 3D matrices were analyzed using a non-destructive contactless rheometer. We observed that the elastic modulus of collagen matrices significantly increased when crosslinked with EDC or functionalized with different molecular weights of HA (**Figure 1E**). We also observed a slight, albeit insignificant, increase in the elastic modulus from crosslinked collagen (209.9 +/- 22.9 Pa) to LMW-HA (216.9 +/- 17.4 Pa), MMW-HA (231.3 +/- 22.1 Pa), and HMW-HA (246.7 +/- 24.9 Pa) matrices, respectively.

Overall, a well-defined collagen matrix system that accurately mimics the native tissue microenvironment of DCs was established, enabling us to investigate the effect of HA molecular weight alone, decoupled from other biophysical parameters, on DC immune potency.

### 3.2. Collagen and HA of different MW regulate availability of cytokines

Cytokine signaling plays a significant role in the immune response of dendritic cells (DCs) and is involved in the regulation of their activation, maturation, migration, and antigen presentation[33,34]. Cytokines exert their effects by binding to specific receptors, thereby triggering intracellular signaling events that regulate their functions in cells[35,36]. Additionally, cytokines are known to bind to ECM components, which in turn control their availability and activity by potentially altering their binding to cell surface receptors[37–39]. To investigate the specific binding of cytokines to collagen and different HA MW, we comprehensively quantified the amount of CCL2, CXCL8, CXCL10, IFN-γ, IL-1β, IL-10, IL-12p70, IL-17A, IL-2, IL-4, IL-6, and TGF-β1 bound to reconstituted matrices using multiplex bead-based ELISA.

As depicted in **Figure 2A and 2B**, we observed a significantly higher amount of CCL2, IFN-γ, IL-1β, IL-6, IL-12p70, and slightly higher amounts of CXCL8 and IL-2 binding to the collagen matrix compared to all other matrix conditions. Previous studies have reported that IL-2, an important stimulator of T cell proliferation, exhibits high binding affinity for collagen type I[39]. However, in the crosslinked collagen matrix, we observed significantly reduced binding for all the cytokines investigated. This reduction in binding could be attributed to the overall reduction in charge of the collagen molecules resulting from the binding of EDC to the amino and carboxyl groups on collagen during crosslinking[40].

**Figure 2:**
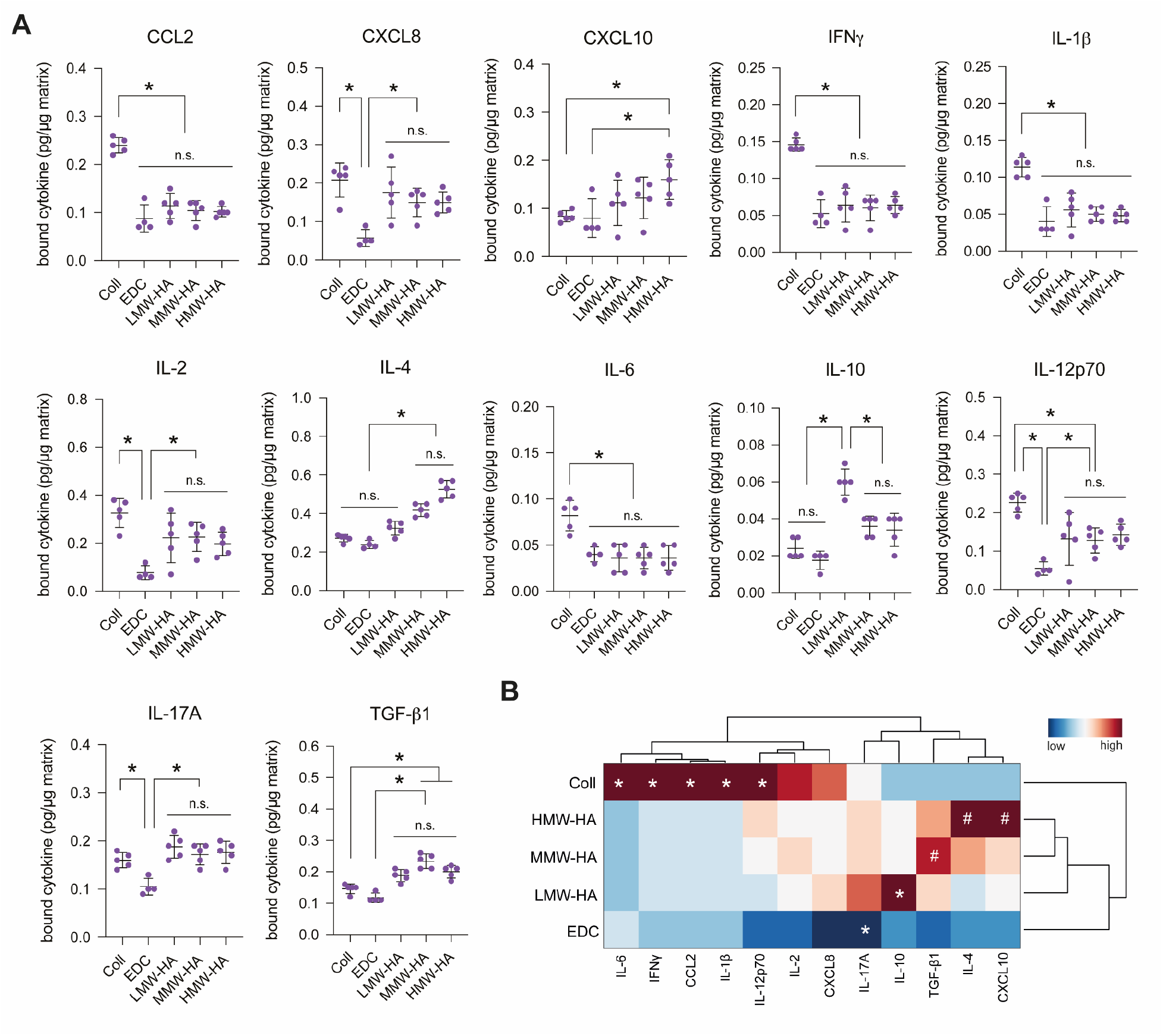
Quantitative analysis of cytokine binding to collagen and different HA MW matrices. Matrices were incubated with 10 µg/mL of cytokines overnight. **(A)** The amount of cytokines bound to the matrices was quantified in picograms per microgram of the matrix (n=6). The significance levels are denoted by * for p ≤ 0.05 and n.s. for non-significant, determined using the Mann–Whitney test. **(B)** Heatmap representing the amount of each cytokine bound to each matrix condition. In the heatmap, asterisks (*) indicates significance compared to all conditions, and hash (#) indicates significance compared to collagen alone.

When HA was present in the matrix, we observed distinct patterns of cytokine binding compared to collagen and crosslinked collagen conditions. Interestingly, the binding characteristics of different cytokines to collagen-HA matrices depended on the MW of HA. We found that CCL10 and IL-4 exhibited significantly high binding to HMW-HA matrices, with the binding decreasing in MMW-HA matrices and further decreasing in LMW-HA matrices. On the other hand, IL-10 showed significantly higher binding to LMW-HA matrices compared to all other matrix conditions. These findings may suggest an additional mechanism through which LMW-HA in the ECM enhances pro-inflammatory immune responses by reducing the availability of anti-inflammatory cytokines such as IL-10. Furthermore, TGF-β1 displayed significantly increased binding to HA-bound matrices compared to collagen matrices. This finding aligns well with a previous report that utilized HA-conjugated hydrogel for the sustained release of TGF-β1[40].

In summary, our study demonstrated that the presence or absence of HA, as well as its molecular weight of HA, can modulate cytokine availability, highlighting the role of the ECM and its composition as a reservoir for cytokines in shaping local tissue functions. It is important to note that our study did not investigate whether the bound cytokines are stably attached to the matrix or their release dynamics over time, if any.

### 3.3. LMW-HA downregulates CD44 receptor expression and upregulates CD11c on iDCs

To study the molecular weight-dependent effect of HA on immature dendritic cells (iDCs), THP-1 cells were differentiated into iDCs in the presence of GM-CSF and IL-4 for 3 days on reconstituted matrices with and without HA presence. Initially, we visualized the cells upon differentiation into iDCs and observed a significant increase in the formation of iDC clusters in samples cultured on MMW-HA and HMW-HA, compared to LMW-HA and collagen matrices (**Figure 3A and 3B**). Furthermore, iDCs cultured in LMW-HA and MMW-HA appeared to have more elongated dendrites (**Figure 3A**), and analysis of cell morphology revealed an increase in the number of spread cells (aspect ratio > 1.5) compared to other matrix conditions (**Figure 3C**). In contrast, iDCs cultured in collagen, crosslinked collagen, and HMW-HA matrices exhibited a low number of clusters (**Figure 3B**) and a more rounded morphology (**Figure 3C**).

**Figure 3:**
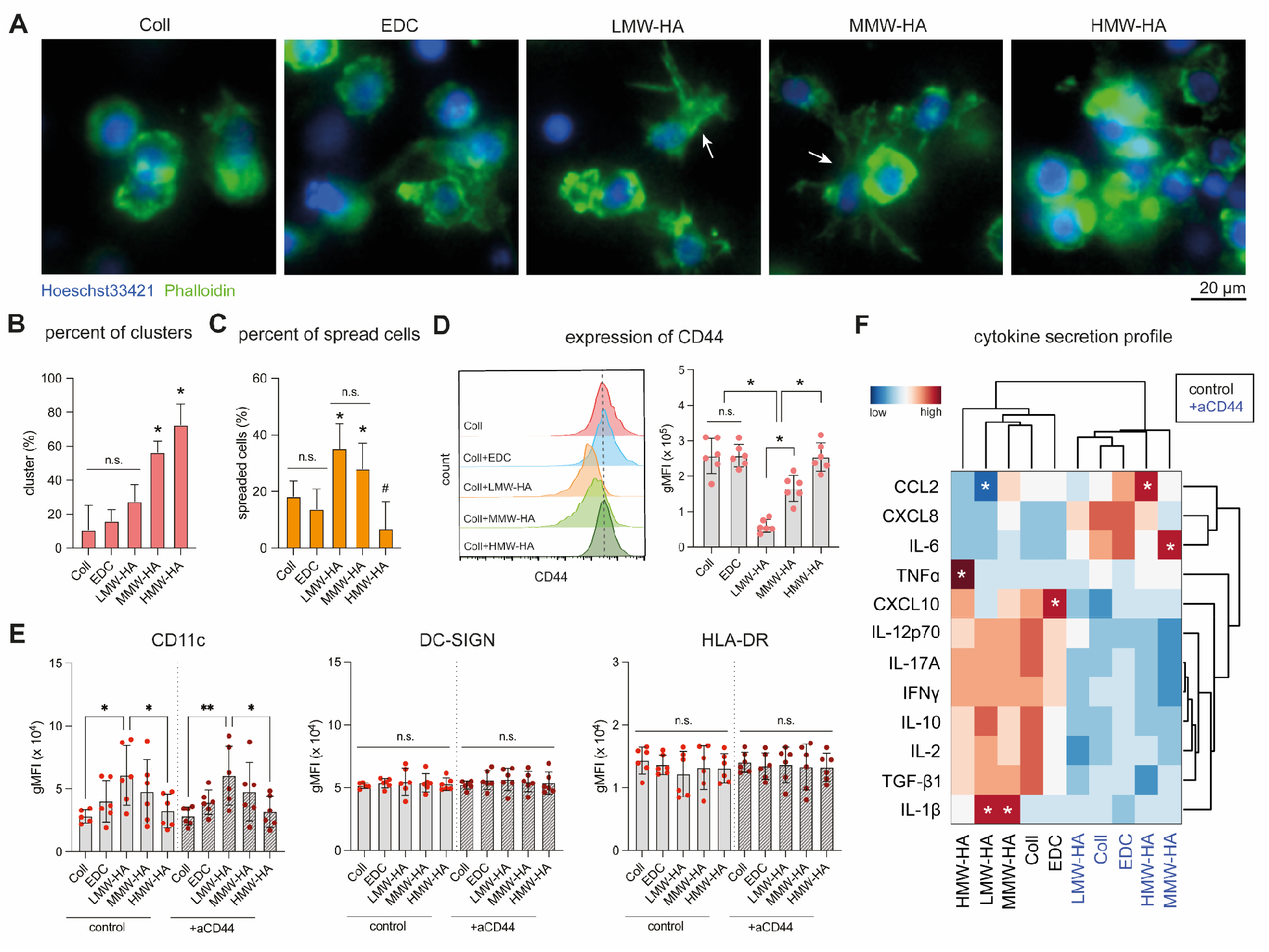
Quantitative analysis of surface marker expression and cytokine secretion profiles of iDCs in Collagen-HA matrices. Human THP-1 monocytes were differentiated into iDC in reconstituted matrices for 3 days. **(A)** Representative images of iDCs within each of the studied matrix conditions. **(B)** Quantitative analysis of the aspect ratio percent of iDC clusters for each matrix condition. **(C)** Quantitative analysis of the percentage of spread iDCs for each matrix condition. **(D)** Analysis of the geometric mean fluorescence intensity (gMFI) for the expression of the CD44 receptor on iDCs. **(E)** Analysis of the gMFI for the expression of surface markers associated with iDC characteristics, namely CD11c, DC-SIGN and HLA-DR. **(F)** Heatmap illustrating the cytokine secretion profile of iDCs in each matrix condition. Experiments were performed with at least 6 replicates. Data are presented as mean ± SD. Asterisks (*) and "n.s." indicate a significance level of p ≤ 0.05 and non-significance, respectively, as determined by the Mann–Whitney test.

Next, we investigated the expression of CD44, one of the key receptors that interacts with HA, using flow cytometry. As shown in **Figure 3D**, CD44 expression was significantly reduced in the presence of LMW-HA and MMW-HA, while HMW-HA exhibited similar levels of CD44 expression compared to collagen and crosslinked collagen matrices.

We further investigated DC differentiation by examining the expression of major DC surface markers, namely CD11c, DC-SIGN (CD209), and HLA-DR. Considering the importance of CD44 as an HA receptor and the observed decrease in CD44 expression in LMW-HA and MMW-HA, we blocked the CD44 receptor by incubating cells with anti-CD44 antibody prior to DC differentiation and supplemented the differentiation media with anti-CD44 antibody. As shown in **Figure 3E**, significant changes were observed in the expression of CD11c, but not DC-SIGN and HLA-DR. The expression levels of CD11c appeared to increase as HA molecular weight decreased from HMW-HA to MMW-HA and LMW-HA, and CD11c expression in HMW-HA was similar to that in iDCs on collagen matrices with and without EDC crosslinking. CD11c is an integrin alpha X chain protein widely used as a differentiation marker for dendritic cells[41]. The increased expression of CD11c in LMW-HA matrices indicates that during pathological conditions where HMW-HA is broken down, the produced LMW-HA contributes to the enhancement of DC differentiation in such tissues. Blocking the CD44 receptor resulted in a similar trend of CD11c expression in iDCs, suggesting that the expression of CD11c was independent of the CD44 receptor (**Figure 3E**). The expression of DC-SIGN and HLA-DR also appeared to be independent of the CD44 receptor.

In addition to characterizing iDCs based on the expression of specific cell surface markers, the secretion of cytokines is also a critical functional response of iDCs. We evaluated the cytokine secretion profiles of iDCs using a multiplex bead-based ELISA. The heatmap in **Figure 3F** shows that iDCs secrete distinct cytokines in different matrix conditions. Our data indicates a generally increased secretion of CXCL10, IFN-γ, IL-1β, IL-2, IL-10, IL-12p70, IL-17A, and TGF-β1, and reduced secretion of CCL2, CXCL8, and IL-6 for all the matrix conditions investigated. However, this trend is reversed when the CD44 receptor is blocked (see Figure 3E), highlighting the importance of CD44 in modulating iDC functions. Additionally, there is a notably increased secretion of IL-1β in both the LMW-HA and MMW-HA matrices (**Figure 3F**). IL-1β secretion by dendritic cells is important for the development of proinflammatory immune responses[42,43]. Therefore, we hypothesize that this may indicate that MMW-HA and LMW-HA contribute to enhancing the proinflammatory immune response of iDCs. This increased IL-1β secretion is consistent with a previous study that found an increased secretion of IL-1β in DCs treated with LMW-HA[30]. However, contrary to this previous study, we observed low secretion of TNF-α by iDCs on LMW-HA matrices and increased secretion of TNF-α on HMW-HA matrices[30]. These differences may be attributed to variations in cell culture dimensionality, as the mentioned study was performed on 2D cell culture plates, whereas our work involves DCs cultured in 3D collagen matrices that better mimic the in vivo tissue microenvironment. We have previously demonstrated that cell culture dimensionality and the conceptual perspective of experimental design are crucial for understanding the immunoregulation of DCs by their microenvironments[8]. Furthermore, the previous study supplemented the cell culture medium with LMW-HA, whereas in our work, LMW-HA is immobilized onto 3D collagen matrices. We also observed a decrease in CXCL10 secretion with decreasing HA MW in the matrices from HMW to MMW and to LMW. CXCL10 is known to be an inflammatory agent and mediates the immune response through its role in the activation and recruitment of T cells to the site of infections[44,45].

Overall, our data suggests that HA fragments enhance the immune potency of iDCs during pathological conditions where HA is broken down. This is supported by the increased expression of the differentiation marker CD11c and enhanced secretion of proinflammatory cytokines, particularly IL-1β. Additionally, we found that the modulation of iDC immune potency by the different MW of HA is independent of the CD44 receptor.

### 3.4. iDC antigen uptake capacity is enhanced by HA fragments via upregulation of CD206 receptor

As previously discussed, the primary role of iDCs is to scavenge the ECM, identify foreign antigens, and engulf them. Therefore, we investigated whether the presence of different MW of HA affects antigen uptake by iDCs. To assess this, iDCs were incubated with FITC-labeled ovalbumin (FITC-OVA), and the uptake was determined by measuring the gMFI of OVA using flow cytometry.

**Figure 4A** illustrates a significant increase in antigen uptake for both LMW-HA and MMW-HA matrix conditions compared to other matrix conditions. This observation is consistent with the trend of increased differentiation, as indicated by CD11c expression (**Figure 3E**). Furthermore, the effect on OVA uptake was found to be independent of the CD44 receptor. Considering that OVA uptake is mediated by the mannose receptor 1 (MMR-1, also known as CD206)[46–48], we proceeded to investigate the expression of the MMR-1 receptor. Our findings revealed a significant increase in MMR-1 expression under LMW-HA and MMW-HA matrix conditions, which appeared to be independent of the CD44 receptor (**Figure 4B**). Moreover, the expression of MMR-1 correlated well with the uptake of OVA (**Figure 4B**).

**Figure 4:**
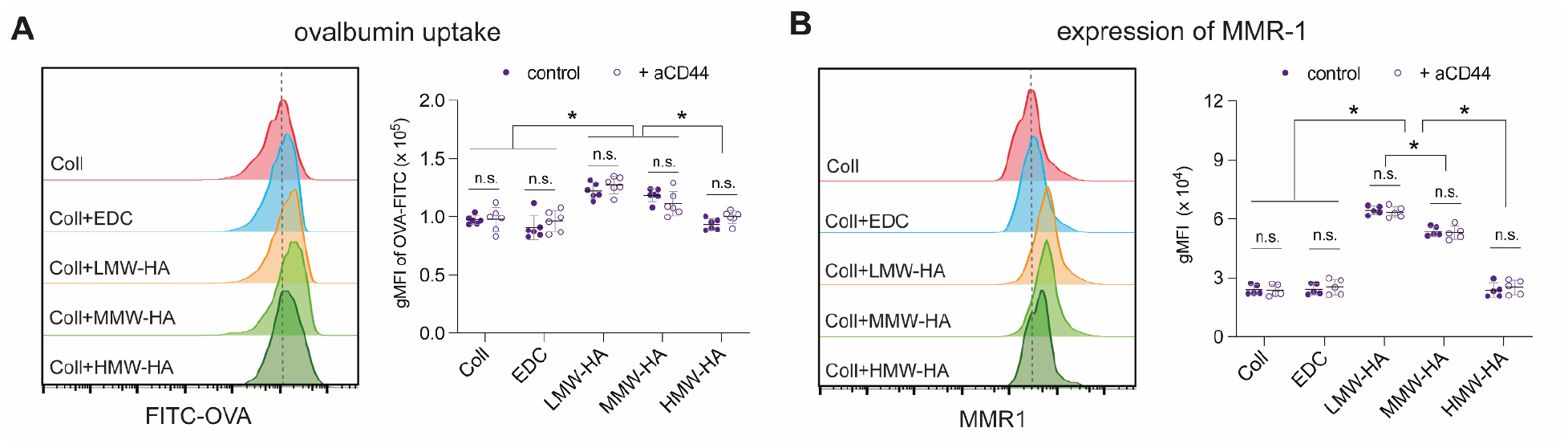
Quantitative analysis of antigen uptake by iDCs. FITC-labeled ovalbumin (FITC-OVA) was added to iDC cultured under each matrix condition for 1 hour. The cells were subsequently analyzed using a flow cytometer to assess their uptake of OVA and MMR-1 receptors, a surface receptor associated with OVA uptake. Representative histogram from flow cytometry analysis illustrating fluorescence intensity and providing quantitative analysis of the geometric mean fluorescence intensity (gMFI) for **(A)** FITC-OVA uptake and **(B)** the expression of MMR-1 receptor. Experiments were performed with a minimum of 5 replicates. Data are presented as the mean ± SD. Asterisks (*) indicates significance at p ≤ 0.05, as determined by the Mann–Whitney test

Taken together, our data indicate that the molecular weight of HA in the ECM has a direct influence on iDC function, specifically their ability to scavenge and uptake antigens. The enhanced antigen uptake observed in the presence of LMW-HA and MMW-HA matrices, along with the upregulation of MMR-1, suggests that targeting HA fragmentation could be a promising approach for modulating antigen presentation and immune responses mediated by iDCs.

### 3.5. Lower molecular weight HA attenuates cytokine secretion of mDCs

iDCs mature into mDCs, which exhibit higher expression levels of HLA-DR, CD80, and CD86[4]. To comprehensively understand the impact of HA with varying molecular weights (MW) on DCs, we also investigated the immunomodulatory potential of mDCs cultured in the different matrix conditions. For this purpose, human monocytic THP-1 cells were differentiated into mDCs using GM-CSF, IL-4, TNFα, and ionomycin for a period of 3 days. Previously, we have demonstrated the effectiveness of this differentiation protocol in generating mDCs with increased levels of costimulatory molecules, MHC proteins, and upregulated immune response pathway genes compared to iDCs, as evidenced by surface marker analysis, cytokine secretion, and transcriptomic profiling[49].

As depicted in **Figure 5A**, we visually observed that mDCs cultured in HMW-HA matrices exhibited a significantly higher formation of cell clusters compared to MMW-HA and LMW-HA matrices. Additionally, we noticed a substantial increase in the formation of dendritic protrusions in MMW-HA and LMW-HA matrices (**Figure 5A**). To validate our visual observations, we quantified the percentage of clusters and spread cells for each matrix condition, confirming our findings (**Figure 5B and 5C**). Subsequently, we examined the expression of the CD44 receptor on mDCs and found no significant change in its expression across all matrix conditions (**Figure 5D**). However, the expression of CD44 on mDCs was significantly lower compared to iDCs in all matrix conditions (**Supplementary Figure S2**). This downregulation of CD44 receptor on mDCs might indicate a mechanism by which mDC enhance their migration towards lymphatic vessels as CD44 receptor deficient bone marrow derived DCs (BMDCs) have been shown to exhibit relatively longer migratory paths compared wild-type BMDCs[21]. Furthermore, we analyzed the surface marker expression specific to mDC differentiation, namely CCR7, HLA-DR, and CD86, both in the presence and absence of anti-CD44 antibodies (**Figure 5E**). CCR7 is involved in the migration of mDCs to lymphoid organs through interaction with the CCL19 chemokine[50,51]. HLA-DR and CD86 are important surface markers that interact with the T cell receptor and costimulatory receptors, respectively, during T cell activation[52,53]. Thus, the expression levels of HLA-DR and CD86 on mDCs reflect the stability of their interaction with T cells and consequently their efficacy in priming T cell response[54]. Surprisingly, our results revealed no significant differences in the surface marker expressions of CCR7, HLA-DR, and CD86 on mDCs across all matrix conditions, regardless of the presence of CD44 (**Figure 5E**). It has been hypothesized that as DCs mature, they become less responsive to external stimuli[49].

**Figure 5:**
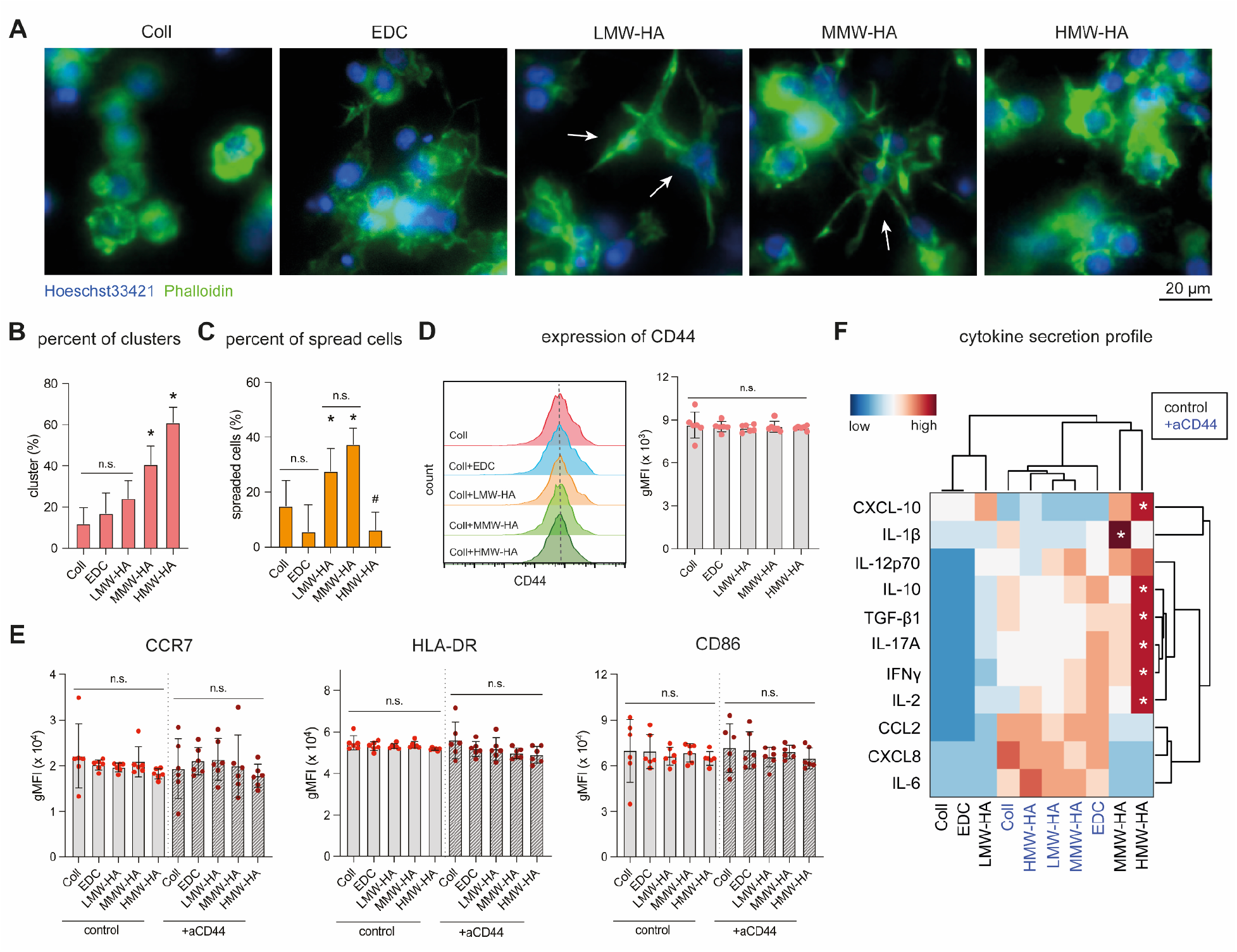
Quantitative analysis of surface marker expression and cytokine secretion profiles of mDCs in Collagen-HA matrices. Human THP-1 monocytes were differentiated and matured into mDCs in reconstituted matrices for 3 days. **(A)** Representative images of mDCs within each of the studied matrix conditions. **(B)** Quantitative analysis of the aspect ratio percent of mDC clusters for each matrix condition. **(C)** Quantitative analysis of the percentage of spread mDCs for each matrix condition. **(D)** Analysis of the geometric mean fluorescence intensity (gMFI) for the expression of the CD44 receptor on mDCs. (E) Analysis of the gMFI for the expression of surface markers associated with mDC characteristics, namely CCR7, HLA-DR and CD86. **(F)** Heatmap illustrating the cytokine secretion profile of mDCs in each matrix condition. Experiments were performed with at least 6 replicates. Data are presented as mean ± SD. Asterisks (*) and "n.s." indicate a significance level of p ≤ 0.05 and non-significance, respectively, as determined by the Mann–Whitney test.

In addition to the analysis of mDC-specific cell surface markers, we examined the cytokine secretion profiles of mDCs in different matrix conditions (**Figure 5F**). We observed that mDCs cultured on HMW-HA and MMW-HA matrices secreted significantly higher amounts of CXCL10, IFNγ, IL-2, IL-10, IL-17A, and TGF-β1 compared to cells cultivated in other matrices, which exhibited a decreased secretion of all cytokines (**Figure 5F**). Similar to the observations in iDCs, CD44 appears to play a role in the extent of cytokine secretion by mDCs, as we measured a general increase in cytokine secretion when CD44 was blocked. Notably, we observed an elevated secretion of IL-6, CXCL8, and CCL2 by mDCs when the CD44 receptor was blocked, consistent with the earlier observations in iDCs.

In summary, our data suggests that although different MWs of HA do not impact the expression of surface markers on mDCs, HA with varying MWs differentially modulates cytokine secretion by mDCs, and this effect is dependent on the CD44 receptor.

### 3.6. MMW-HA and HMW-HA reduced DC migration in CD44 dependent manner

Migration plays a crucial role in the function of DCs, whether it involves iDC scavenging the ECM and uptaking antigens, or mDC transitioning to lymphoid organs to locate and activate lymphocytes, thereby initiating the adaptive immune response[51,55–57]. In this study, we investigated how different HA MW affects the migratory capacity of both iDCs and mDCs and whether this effect is dependent on the CD44 receptor. To accomplish this, z-stack images of both iDCs and mDCs were captured at 5 μm intervals, with a total z-layer thickness of 800 μm, using a fluorescence microscope. The percentage of migrated DCs and the maximum depth of migration into the matrices were quantified using a custom-built image analysis toolbox. A schematic illustration of the cell migration analysis is presented in **Figure 6A**.

**Figure 6:**
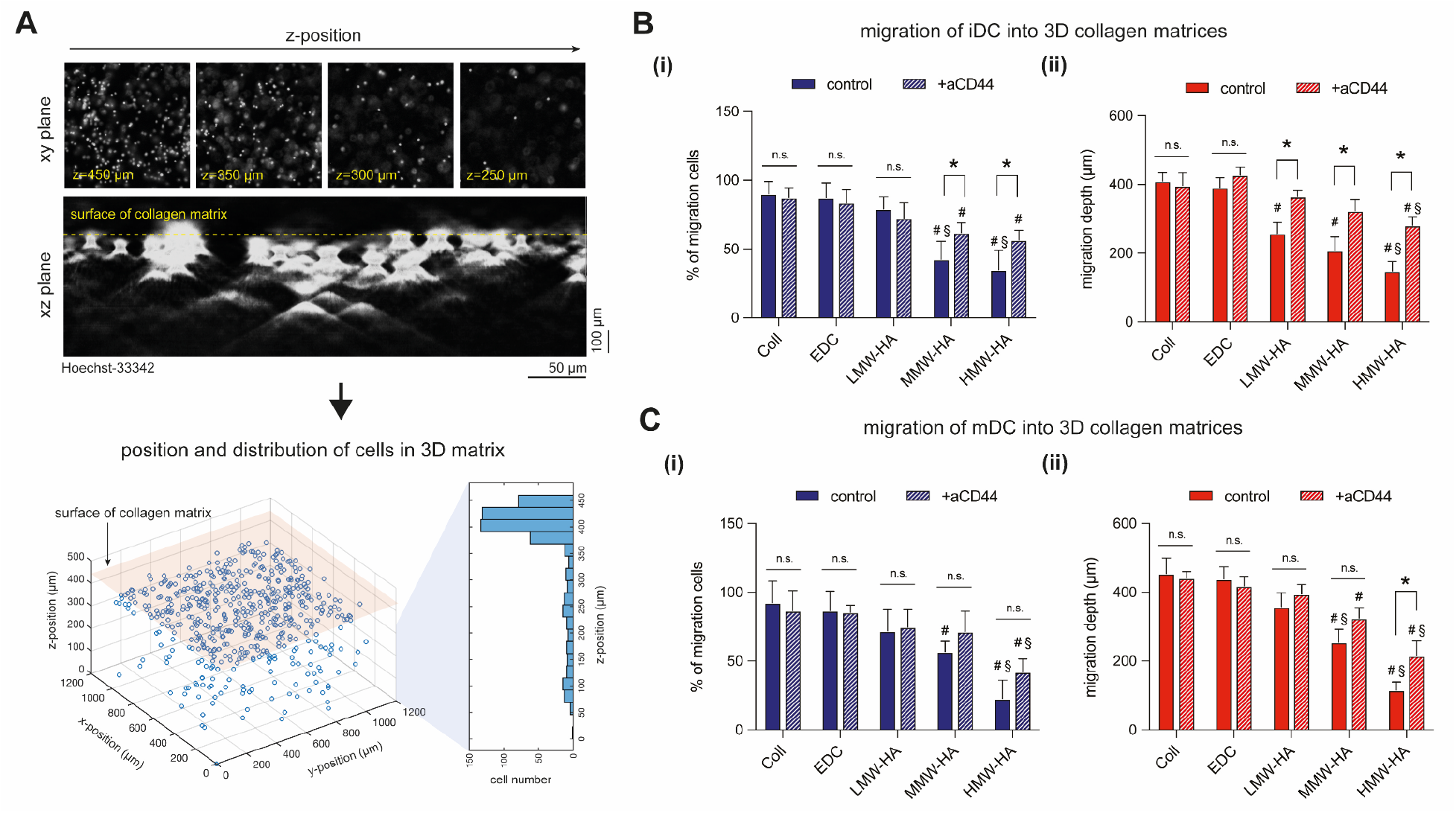
Quantitative Analysis of Migratory Capacity of iDCs and mDCs. (A) Representative images and schematic representation of DCs at different z-positions within the matrices. (B) Quantitative analysis of (i) percent of migrating cells and (ii) depth of migration of iDCs. (C) Quantitative analysis of (i) percent of migrating cells and (ii) depth of migration of mDCs. Data are presented as the mean ± SD. Asterisk (*) indicates significance at p ≤ 0.05. Hash (#) and section symbol (§) indicate significance compared to collagen and LMW-HA matrices, respectively.

For iDCs, no significant difference in the percentage of cells migrating into the collagen, crosslinked collagen, and LMW-HA matrices was observed (**Figure 6B(i)**). However, there appears to be a decrease in the percentage of migrating cells in MMW-HA and HMW-HA matrices. Blocking the CD44 receptor resulted in a significant increase in the percentage of migrating cells in both MMW-HA and HMW-HA matrices, but not in the other matrix conditions. In addition to the percentage of migrating cells, we also quantified the maximum migration depth. We observed a significant reduction in iDC migration in all HA-bound matrices, with a dependence on HA molecular weight. An increase in HA MW was found to decrease the migration of iDCs (**Figure 6B(ii)**). However, upon blocking the CD44 receptor, the migration depth of iDCs significantly increased in LMW-HA, MMW-HA, and HMW-HA matrices, suggesting that HA may provide an adhesive microenvironment for iDCs.

For mDCs, the percentage of migrating cells was significantly reduced in MMW-HA and HMW-HA matrices compared to all other matrices (**Figure 6C(i)**). Blocking the CD44 receptor led to an increase in the depth of migrating cells in both MMW-HA and HMW-HA matrices, with significance observed only in HMW-HA. Similar to the percentage of migrating cells, the maximum migration depth of mDCs was significantly decreased in MMW-HA and HMW-HA matrices (**Figure 6C(ii)**). However, blocking the CD44 receptor enabled mDCs to migrate into MMW-HA and HMW-HA matrices with increased migration depth.

Overall, our data suggest that MMW-HA and HMW-HA, but not LMW-HA, reduce DC migration for both iDCs and mDCs in a CD44-dependent manner. These results also indicate that HA of different molecular weights can provide distinct adhesive microenvironments for DCs, as observed in other cell types[18,58,59]. The regulation of the adhesive microenvironment may support DC functions in specific tissues. For example, since LMW-HA is present in injured tissues, we hypothesize that LMW-HA facilitates the migration of iDCs to scan for antigens and supports the migration of mDCs into nearby lymph nodes. Conversely, MMW-HA and HMW-HA restrict DC migration, allowing them to fulfill their functions, such as antigen presentation to T cells within lymphoid organs. Coincidentally, the migratory reduction due to increase in MW of HA trends with the expression of Cd11c from iDCs. However, the role of integrins such as CD11c in DC and macrophage migration remains controversial[60].

## 4. General discussion and conclusions

HA plays a crucial role in numerous physiological and pathological conditions and exists in both soluble and bound forms within the ECM. Physiologically, HA in the ECM predominantly exists as HMW forms during tissue homeostasis. It has been shown that various tissues possess different average HA molecular weights. For example, lymph nodes have been found to contain HA with a molecular weight of 790 kDa[61]. However, under certain pathological conditions such as tissue injury, inflammation, and tumor development, HA undergoes degradation, leading to the generation of smaller molecular weight fragments. The extent to which the molecular weight of HA contributes to the establishment of a microenvironment that facilitates specific cellular functions remains largely unexplored. In this study, we demonstrate that the molecular weight of HA present in the ECM can influence binding to various cytokines, thereby regulating cytokine availability to cells. We then show that distinct HA molecular weights differentially modulate the immune response of DCs. Specifically, LMW HA enhances the differentiation and antigen uptake of iDCs, while MMW and HMW HA enhance the cytokine secretion of mDCs. Our findings are consistent with a study of HA MW using overexpression of HYAL1 and tamoxifen-inducible HYAL1 expression in mouse skin, which revealed that generating smaller HA fragments can promote DC differentiation, as indicated by high CD11c expression[62]. We also observed that the specific cell surface marker expression of both iDCs and mDCs was independent of the CD44 receptor, but not cytokine secretion profiles. When CD44 receptors were blocked with antibodies, secretion of CCL2, CXCL8, and IL-6 was higher in both DC populations compared to the control. The simultaneous expression of these cytokines has been reported to correlate with acute inflammation[63]. Our data therefore suggests the possible role of CD44 in regulating acute inflammation. While the RHAMM is also an important HA receptor particularly with its involvement in modulation of cell motility[64], it has been suggested that the CD44 function is dominant[65,66]. Hence, this study focused on understanding the CD44 dependence with regards to how different MW of HA affect DCs. We would like to clarify that the measured cytokine secretion profiles discussed here do not take into account the proportion of cytokines bound to the matrix. Further work is needed to understand the bioactivity of matrix-bound cytokines on DC biology and their release dynamics. This is beyond the scope of this study. Besides, the bioavailability of cytokines in our in vitro system is reflective of the collagen-HA portion of the tissue niche. Additionally, we found that CD44 expression is higher in iDCs compared to mDCs. Furthermore, our results demonstrated that MMW-HA and HMW-HA reduced the migratory capacity of both iDCs and mDCs, while blocking CD44 appeared to erase the adhesiveness of HA, enhancing migration in both cell types. These findings are consistent with a study demonstrating that the degradation of HA by HYAL1 can enhance the migration of DCs from the skin[62]. Together it can be interpreted that larger molecular weight HA could present as a more adhesive microenvironment for DCs.

Based on our findings and depicted in **Figure 7**, we propose a physiological model whereby the DC microenvironment is instructional. It demonstrates that iDC functions are enhanced when LMW-HA is present, such as in inflammation, injured tissue, and cancer tissue. In these specific tissues, iDCs increase their uptake and migration. Once activated through uptake, they progressively mature into mDCs. Concurrently, they would have migrated towards the nearest lymphatic tissue or adjacent tissues where MMW- and HMW-HA are present, serving their functions and the need for motility diminishes. Our study sheds light on the significance of HA molecular weight in physiological and pathological processes, highlighting its role in modulating cytokine availability, immune response, and cell migration. Further needed investigation into the impact of HA molecular weight on antigen presentation to T cells and the precise mechanisms underlying HA-CD44 interactions will deepen our understanding of the complex interplay between HA and immune cells.

**Figure 7:**
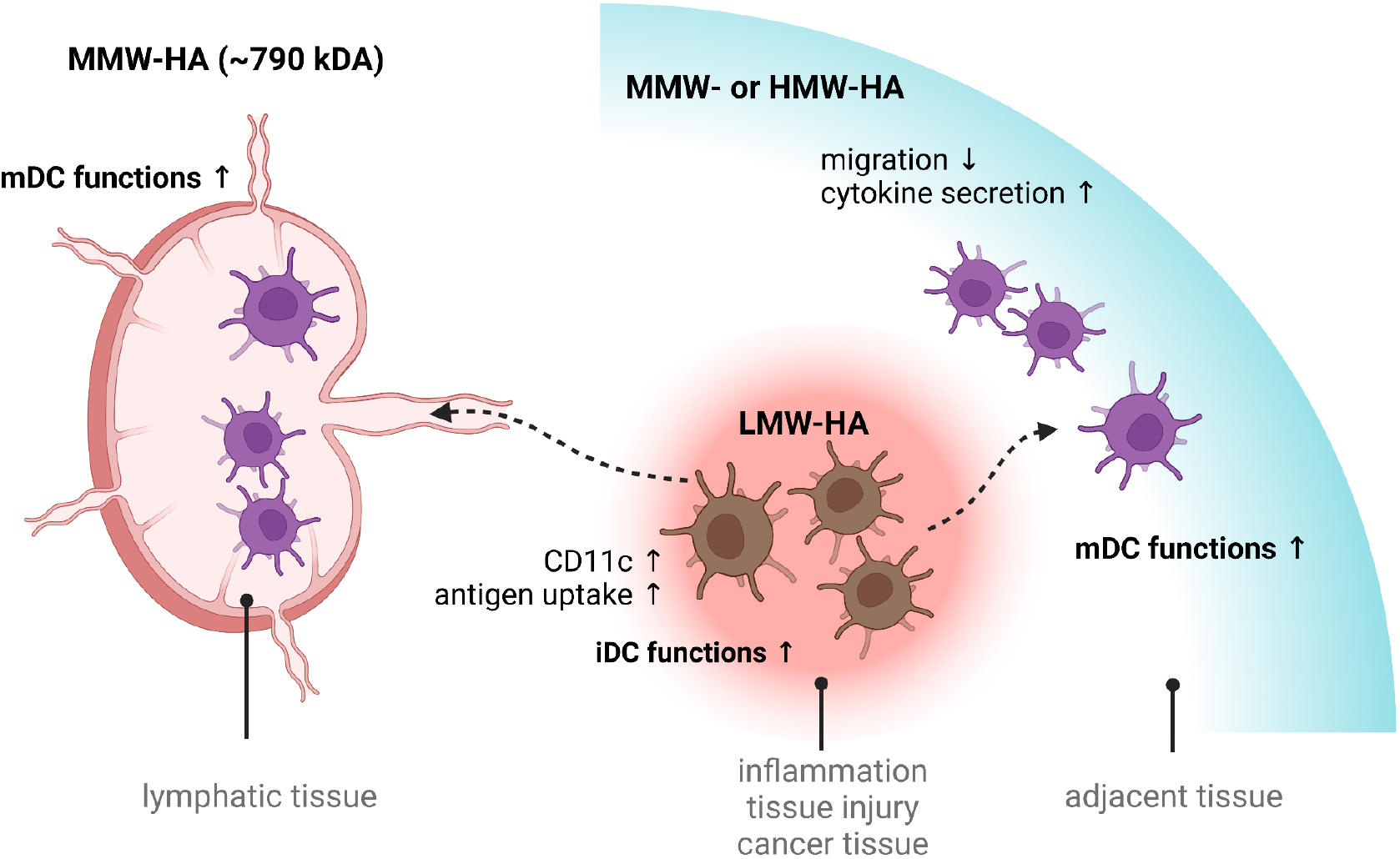
Schematic illustration of proposed HA MW dependent effect on DC immune potency.

Overall, our findings demonstrate that HA molecular weight is capable of shaping adaptive immune responsiveness via dendritic cells, which is relevant during tissue injury, inflammation, and cancer progression. The fundamental knowledge obtained from this study provides valuable insights that can be incorporated into the design of HA-based hydrogels with controlled release properties, the development of DC-based disease modeling (e.g., psoriasis and atopic dermatitis), and therapy. Based on our findings, LMW-HA could be a potential biomolecule that enhances DC differentiation, ultimately increasing the efficiency of DC vaccine generation.

## Data Availability

The datasets generated during and/or analyzed during the current study are available from the corresponding author on reasonable request. Script for topological analysis cab be assessed here: https://git.sc.uni-leipzig.de/pe695hoje/topology-analysis

## Acknowledgement

The authors acknowledge the support from New York University Abu Dhabi (NYUAD) Faculty Research Fund (AD266). The authors would also like to acknowledge support from NYUAD core technology platform. Experiments were performed using the NYUAD Light Microscopy and Molecular and Cell Biology Platforms.

## Competing interests

The authors declare no competing interests

## Notes

### Competing Interest Statement

The authors have declared no competing interest.

