## Supplemental Information for "Tissue-bound hyaluronan molecular weight as a regulator of dendritic cell immune potency"

Prof. Dr. Jeremy Teo

| | |
|---|---|
| Postal address | New York University Abu Dhabi<br>Division of Engineering<br>Abu Dhabi, UAE. |
| Telephone | +971 2 6286689 |
| E-mail: | |

**Supplementary Table 1:** Antibodies used for flow cytometry in this study

| Marker | Fluorochrome | Host/Target | Isotype | Clone | Catalog Number |
|---|---|---|---|---|---|
| CCR7 | Alexa Fluor 488 | Mouse anti-Human | IgG2a, κ | G043H7 | 353206 |
| CD11c | PerCP | Mouse anti-Human | IgG1, κ | Bu15 | 337234 |
| CD44 | Alexa Fluor 594 | Mouse anti-Human | IgG1, κ | C44Mab-5 | 397510 |
| CD86 | Brilliant Violet 605 | Mouse anti-Human | IgG1, κ | BU63 | 374214 |
| CD206 | Brilliant Violet 510 | Mouse anti-Human | IgG1, κ | 15-2 | 321138 |
| CD209 | PE | Mouse anti-Human | IgG2a, κ | 9E9A8 | 330106 |
| HLA-DR | Brilliant Violet 421 | Mouse anti-Human | IgG2a, κ | L243 | 307636 |

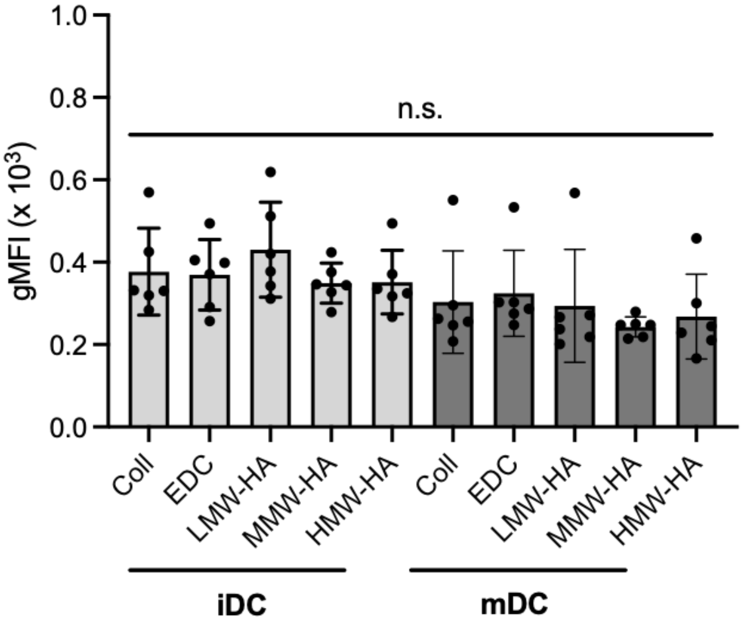


**Supplementary Figure S1: Quantitative analysis CD44 receptor after blocking with anti-CD44 antibody.** THP-1 cells were with 2 µg/ml of anti-CD44 antibodies for 20 minutes prior to iDC and mDC differentiation. Successful blocking was confirmed by staining with anti-human CD44 antibody conjugated with Alexa Fluor 594 and analyzing using flow cytometry. Plot shows quantitative analysis of the gMFI for the presence of CD44 receptor after blocking experiment. Experiments were performed with at least 6 replicates. Data are presented as mean ± SD. Asterisks (*) and "n.s." indicate a significance level of $p \leq 0.05$ and non-significance, respectively, as determined by the Mann–Whitney test.

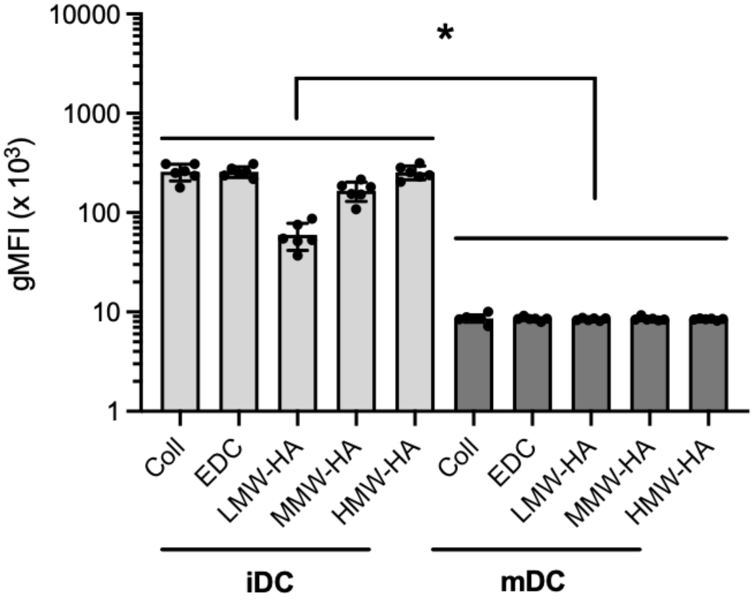


**Supplementary Figure S2: Comparison of CD44 receptor expression between iDCs and mDCs.** Data is replotted from Figure 3D and 5D to emphasize the difference in CD44 receptor expression for iDCs and mDCs. Experiments were performed with at least 6 replicates. Data are presented as mean ± SD. Asterisks (*) and "n.s." indicate a significance level of $p \leq 0.05$ and non-significance, respectively, as determined by the Mann–Whitney test.
